# Deciphering Brain Organoids Heterogeneity by Identifying Key Quality Determinants

**DOI:** 10.1101/2025.01.13.632763

**Authors:** Daniil Kachkin, Naime Zagha, Tom Boerstler, Federica Furlanetto, Negar Nayebzade, Luke Zappia, Michelle Boisvert, Michaela Farrell, Sonja Ploetz, Martin Regensburger, Claudia Günther, Juergen Winkler, Pooja Gupta, Fabian Theis, Marisa Karow, Sven Falk, Beate Winner, Florian Krach

## Abstract

Brain organoids derived from human pluripotent stem cells (hPSCs) hold immense potential for modeling neurodevelopmental processes and disorders. However, their experimental variability and undefined organoid selection criteria for analysis hinder reproducibility. As part of the Bavarian ForInter consortium, we generated 72 brain organoids from distinct hPSC lines. We conducted a comprehensive analysis of their morphological and cellular characteristics at an early stage of their development. In our assessment, the Feret diameter emerged as a reliable, single parameter that characterizes brain organoid quality. Transcriptomic analysis further confirmed the reliability of this marker and identified a negative impact of mesenchymal cells on the abundance of organoid formation. High-quality organoids consistently displayed a lower mesenchymal cell presence. These findings offer a framework for enhancing brain organoid standardization and reproducibility, underscoring the need for morphological quality controls and the consideration of mesenchymal cell influence on organoid-based modeling.

**Graphical Abstract:** 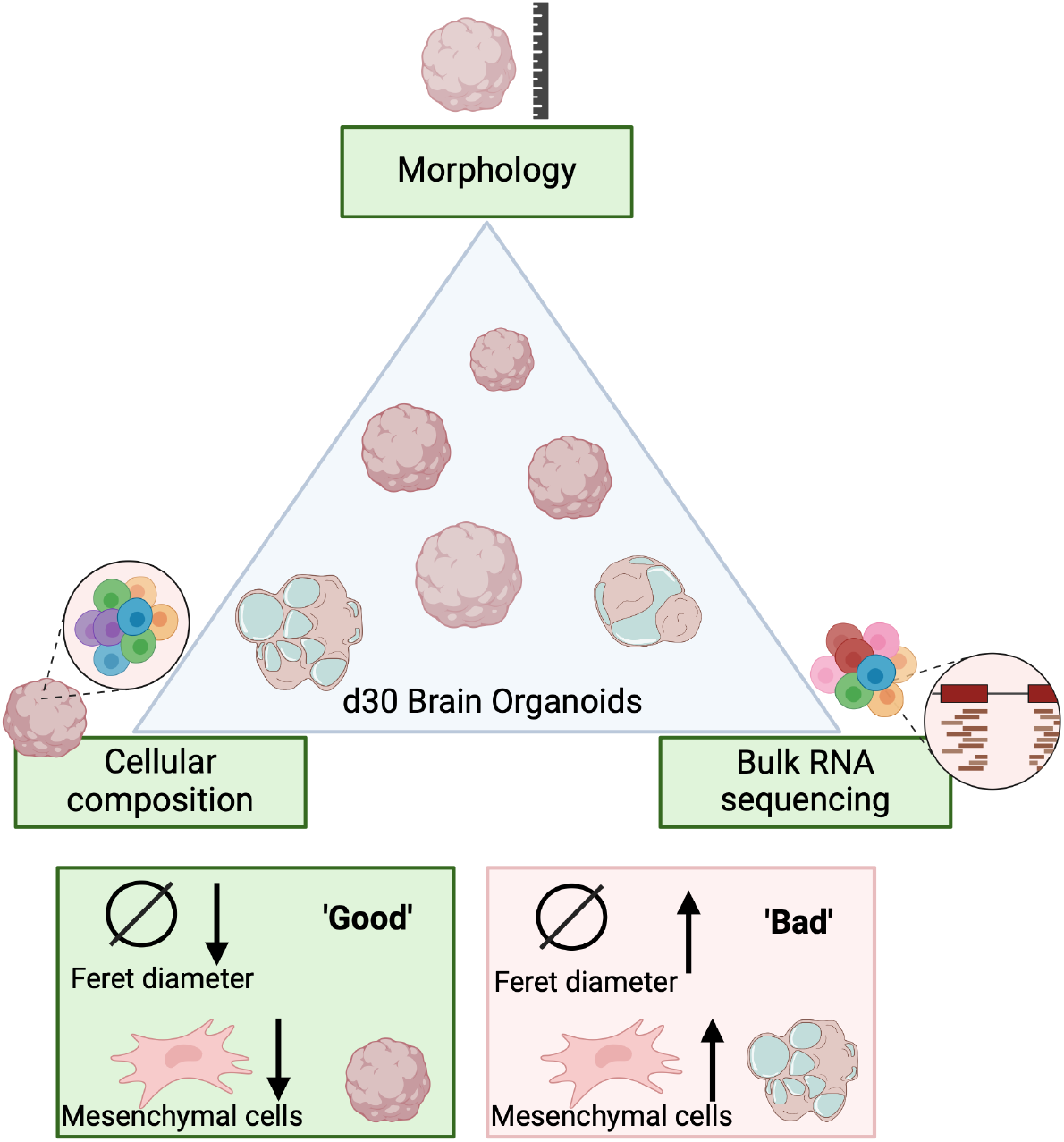

## Introduction

The creation of self-organizing 3D neural structures from human pluripotent stem cells (hPSC) was a milestone in stem cell research^1^. Brain organoids mimic aspects of human brain development, including cellular architecture and diversity. Major advantages of such models include their spatial organization and more physiological cell-cell communication in comparison with 2D cultures^2,3^. Recent discoveries in molecular and cellular anthropology and human brain evolution underline their methodological power^4–8^. Additionally, the system provides a useful platform to model a range of neurological disorders. These include rare developmental disorders, such as Optiz syndrome^9^ or microcephaly^10,11^, neurodegenerative disorders like Alzheimer’s disease^12,13^, or amyotrophic lateral sclerosis (ALS) ^14^, and infectious diseases like Zika virus (ZIKV) infection^15,16^.

Despite these remarkable advances, challenges related to robustness, accuracy, and reproducibility still exist. Hence, organoid models should be evaluated carefully^17^. Brain organoids produced from the same cell line, even under identical conditions, show variations in spatial organization and cellular diversity^18^. Publications often fail to provide clear criteria for selecting organoids for experiments. They typically do not describe if and how many organoids were discarded during the differentiation process^19–21^. This creates challenges in assessing the reliability of experimental results and introduces potential bias in the selection of organoids for disease modeling. While in principle, any human stem cell line can be used to generate brain organoids, only a few specific cell lines have been used (e.g., hESC lines H9 (WA09), H1 (WA01), and WIBR1/2/3 and iPSC lines IMR90 and Kucg2)^22–24^. However, international guidelines recommend using a diverse array of human stem cell lines to ensure the robustness and generalization of findings^25^.

Here, we aim to understand early determinants of brain organoid quality by generating a set of brain organoids from 12 different hPSC lines using an adaptation of the original Lancaster protocol^19^. All cell lines were obtained from healthy donors. After 30 days of differentiation, significant variability was observed both within the brain organoids from different PSC origins and among those derived from the same cell line. Through a systematic framework of analyzing morphological features and transcriptional signatures, we identified the organoid Feret diameter as a determinant of organoid quality. Via computational analysis of transcriptomic data, we estimated the cellular composition of our organoids and found a correlation between organoid quality and the proportion of mesenchymal cells. The proportion of mesenchymal cells was also positively correlated with the organoids’ Feret diameter. These findings can serve as a valuable resource for researchers working with brain organoids to handle diversity and reduce variability.

## Results

We selected a panel of 12 hPSC lines consisting of two embryonic stem cell (hESC) lines (H9 and HuES6) and ten iPSC lines, including both commercially available and in-house lines (Table S1). All iPSC lines were analyzed for TRA-1-60 expression as a surrogate marker of pluripotency. All lines had greater than 90% TRA-1-60 positive cells. For brain organoid differentiation, we utilized an optimized version of the original differentiation protocol developed by Lancaster and Knoblich (2014)^19^.

### Cellular and Structural Composition of Brain Organoids

After 30 days of differentiation, we acquired a set of morphologically diverse organoids and characterized their cellular composition and architecture (Fig. 1A, B). We randomly selected six organoids from each line for immunostaining analysis. Cryosections of the organoids were stained with antibodies against MAP2, a neuronal cytoskeleton component, and SOX2, a transcription factor expressed in neural stem cells of the CNS. This allowed us to examine the cellular composition and distribution within the organoids and the formation of ventricular-like structures (VLS) (Fig. 1C). VLSs resemble the ventricular zone (VZ) in the developing brain composed of neural stem cells and hence also the primary site of neurogenesis. Their presence in brain organoids demonstrates tissue organization similar to that of the developing brain, which is important for developmental processes such as the self-renewal of neural stem cells. This analysis revealed considerable variability among the organoids: some organoids failed to form VLS while others developed multiple VLS populated with SOX2+ and surrounded with MAP2+ cells, indicating a high degree of active neurogenesis (Fig. 1D). We randomly selected three organoids from each line and determined the quantities of PAX6- and MAP2-expressing cells. We observed a high degree of variability, both when comparing hPSC lines originating from different donors and among individual organoids originating from a single donor hPSC line (Fig. 1E, F). These findings highlight the heterogeneity in organoid development regarding VLS development and cellular composition.

**Figure 1.**
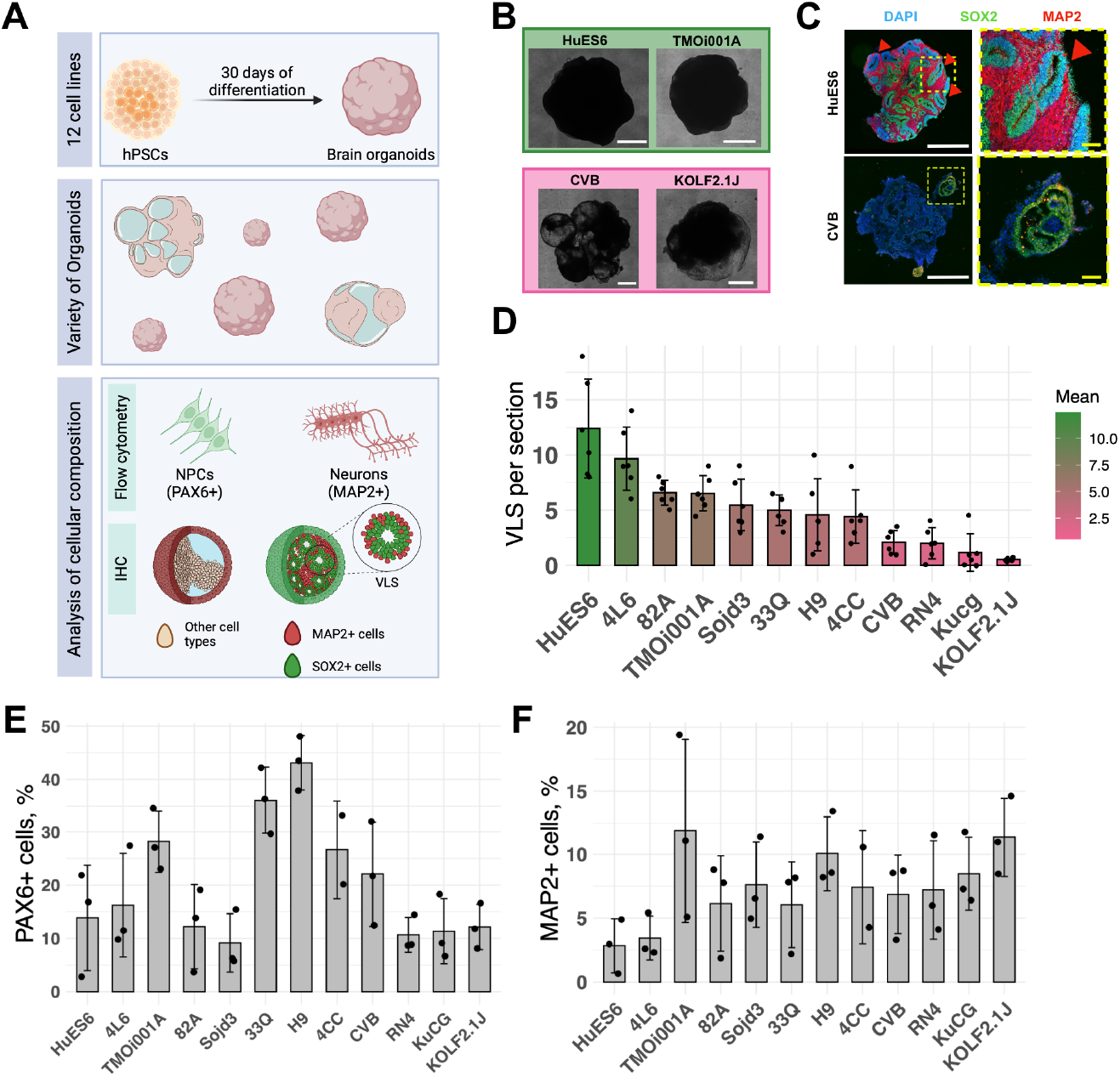
Cellular and structural composition of brain organoids. **A**. Experimental paradigm depicting analysis strategy of 12 hPSC lines at day 30 of brain organoid development for studying morphological diversity, cellular composition (neuronal progenitor cells (NPCs) and neurons) and ventricular-like structure (VLS) formation. **B**. Bright field images of organoids at day 30 of differentiation of the lines HuES6, TMOi001A, CVB, and KOLF2.1J. Scale bar: 1 mm. **C**. Immunofluorescence (IF) images of organoids stained at day 30 with DAPI, SOX2 (green), and MAP2 (red). White scale bar: 1 mm; yellow scale bar: 100 µm. Ventricular-like structures are highlighted with red arrows. **D**. Bar blot depicting the number of VLS per section. Each dot represents the amount of VLS per section of the organoid. The color scale depicts the increase in VLS ranging from green (high amount) to pink (low amount). Data represented as mean ± SD (n = 6 organoids). **E** and **F**. Bar blots with the percentage of PAX6+ (E) or MAP2+ (F) cells of individual organoids assessed by flow cytometry. Data represented as mean ± SD.

### Morphology evaluation of brain organoids

To formalize the expert evaluation commonly used in the field for selecting higher-quality organoids, we employed an unbiased and streamlined analysis approach. We randomly selected 72 individual brain organoids (six per line) for further analysis. Prior to harvesting, an expert with more than 5 years of experience visually evaluated and rated the organoids’ quality as either ‘good’ or ‘bad’. We then captured bright-field images of the organoids and used ImageJ software to measure various morphological parameters. Subsequently, we applied a sequence of statistical analyses to identify the morphological parameters that could predict the expert’s evaluation (Fig. 2A).

**Figure 2.**
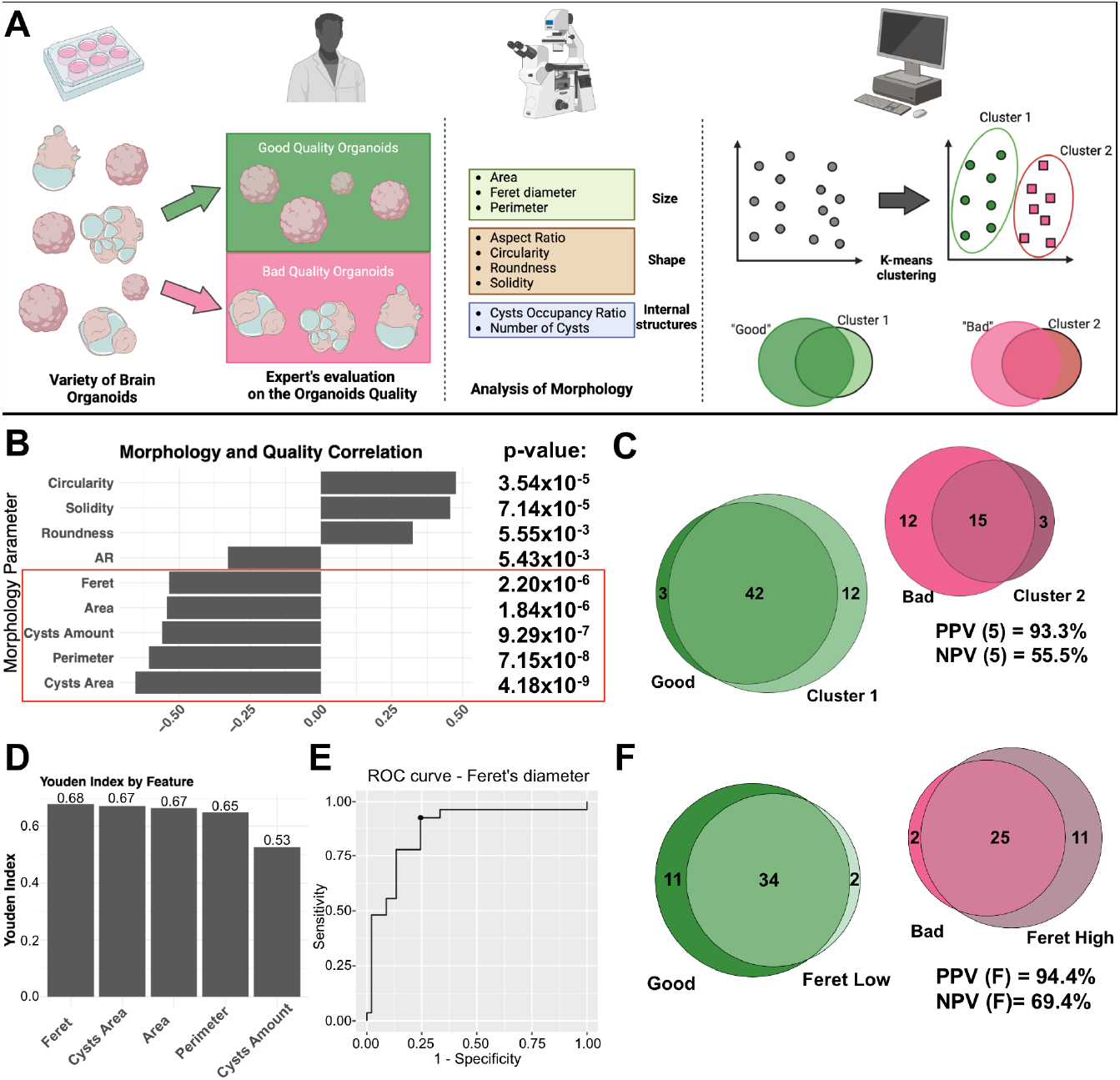
Morphology assessment of brain organoids identifies Feret diameter as a classifier of organoid quality. **A**. Schematic illustrating the strategy to assess and determine morphological parameters that align with expert evaluation. **B**. Bar graph depicting the result of the correlation analysis of morphological features with the expert’s quality evaluation. The red square highlights five parameters surpassing the stringency threshold of p-value < 10^−5^. **C**. Venn diagram depicting the overlap of organoids evaluated by an expert (green and pink, ‘good, ‘bad’’) or clustered in an unbiased way into two groups with k-means clustering using five highly correlated parameters (light-green and dark-pink, ‘Cluster 1’ and ‘Cluster 2’). Positive (PPV) and negative prediction values (NPV) of the k-means clustering are depicted in the graph. **D**. Bar graph illustrating Youden indices showing the diagnostic properties of five morphological parameters. The number on top of the graph represents the index value. **E**. ROC curve for Feret diameter illustrating the ideal threshold with maximized specificity and sensitivity. Identified Feret diameter threshold: 3050 μm. **F**. Venn diagram depicting the overlap of organoids evaluated by an expert (green and pink, ‘good, ‘bad’’) or clustered in two groups using Feret diameter threshold (light-green and dark-pink, Feret Low - Low Feret diameter 1’ and Feret High - High Feret diameter 2’). Positive (PPV) and negative prediction values (NPV) of the k-means clustering are depicted in the graph.

To determine which distinct morphological parameters at day 30 are associated with the expert’s evaluation, we conducted a point-biserial correlation analysis of the nine parameters. We set an FDR-corrected p-value cutoff of 10^−5^ and a point-biserial correlation coefficient |r-value| of > 0.5 as predefined significance thresholds. Five morphological parameters satisfied this threshold: Feret diameter, Area, Cysts Amount, Perimeter, and Cysts Area (Fig. 2B).

To validate our findings, we asked whether these five parameters together can adequately reflect organoid quality in an unsupervised manner using k-means clustering. K=2 was computed as the ideal number of clusters for the given dataset (Supplementary Figure S1). Interestingly, the clusters identified by k-means reflect the experts’ evaluation to a high degree with a positive and negative predictive value (PPV and NPV) of 93.3% and 55.5%, respectively (Fig. 2C). This confirms that morphological measurements can objectify the visual expert evaluation. Next, we investigated if we could simplify this decision-making process to a single morphological parameter. We applied Youden’s J statistics to the five morphological parameters to determine the threshold of a single parameter with the best diagnostic properties. All parameters computed thresholds with Youden indices above 0.5, indicating their positive prognostic properties (Fig. 2D and E). The Feret diameter (a maximal caliper diameter: the longest distance between any two points of the organoid) exhibited the best performance (Youden index of 0.68) at a threshold of 3050 μm (Supplementary Table S2). Strikingly, classifying the organoids by using this single parameter accurately reflected the expert’s evaluation with a PPV and NPV of 94.4% and 69.4%, respectively (Fig. 2F). The similarity of these clustering methods to the expert’s evaluation was also visually evident when putting the originally determined nine morphological parameters into principal component space (Supplementary Fig. S2A-C).

### The fraction of mesenchymal cells affects the morphological quality of brain organoids

To uncover the cellular and molecular basis of organoid quality, we subjected the previously analyzed 72 organoids (Fig. 2) to bulk RNA sequencing for gene expression and cellular deconvolution analysis (Fig. 3A). First, we analyzed the dataset to identify genes that were differentially expressed when classifying the organoids by either the expert’s evaluation, clusters obtained via k-means using five parameters (Fig. 2B) or only the Feret diameter threshold. Differential expression analysis, based on the expert evaluation, revealed the highest number of differentially expressed genes (2647). The other two classifying methods resulted in fewer differentially expressed genes (k-means: 70; Feret diameter: 441), but exhibited a large overlap with the expert’s evaluation, validating our previous analyses. 42 genes were common to all classification methods (Fig. 3B). We performed Gene Ontology (GO) analysis on those 42 genes to uncover the underlying biology behind this observation. The resulting GO analysis suggests the presence of cells not associated with neural lineage. (Fig. 3C). Therefore, we sought to estimate the cellular composition of our organoids and employed BayesPrism deconvolution analysis. BayesPrism is a computational tool that estimates the cellular composition of bulk RNA sequencing data by using a reference single-cell RNA sequencing dataset^26^. As a reference, we used a subset of the Human Neural Organoid Cell Atlas (HNOCA)^27^. The analysis revealed a heterogeneous, mostly neural cellular composition (range: 25.93% - 99,46%). Importantly, we observed a significant variation in the proportion of mesenchymal cells (MC) across the samples, ranging from 0.5% to 74% (Fig. 3D). The variability in mesenchymal cell composition among organoids derived from a single donor was lower (median coefficient of variation: 50.93%; range of coefficient of variation: 5.57% - 105.87%) than the variability observed when comparing the mean mesenchymal cell composition of organoids from different donors (coefficient of variation of mean MC composition in all cell lines: 80.98%). This suggests that mesenchymal cells preferentially arise in specific hPSC lines while acknowledging a certain degree of heterogeneity when comparing organoids of the same donor. The proportion of mesenchymal cells exhibited a significant correlation with the expert evaluation of organoid quality (r = −0.79; p = 2×10^−16^). A significant negative correlation was also evident for the Feret diameter (r = 0.59; p = 5.05×10^−8^), indicating that high-quality organoids are smaller and exhibit a lower content of mesenchymal cells (Fig. 3E-F). We validated this finding computationally by applying WebCSEA - Web-based Cell-type Specific Enrichment Analysis of Genes^28^ to the 42 commonly differentially expressed genes. We found an association of our set of genes with non-neural cells like epithelial and stromal cells (fibroblasts and mesenchymal stem cells) in the dataset (Supplementary Figure S3).

**Figure 3.**
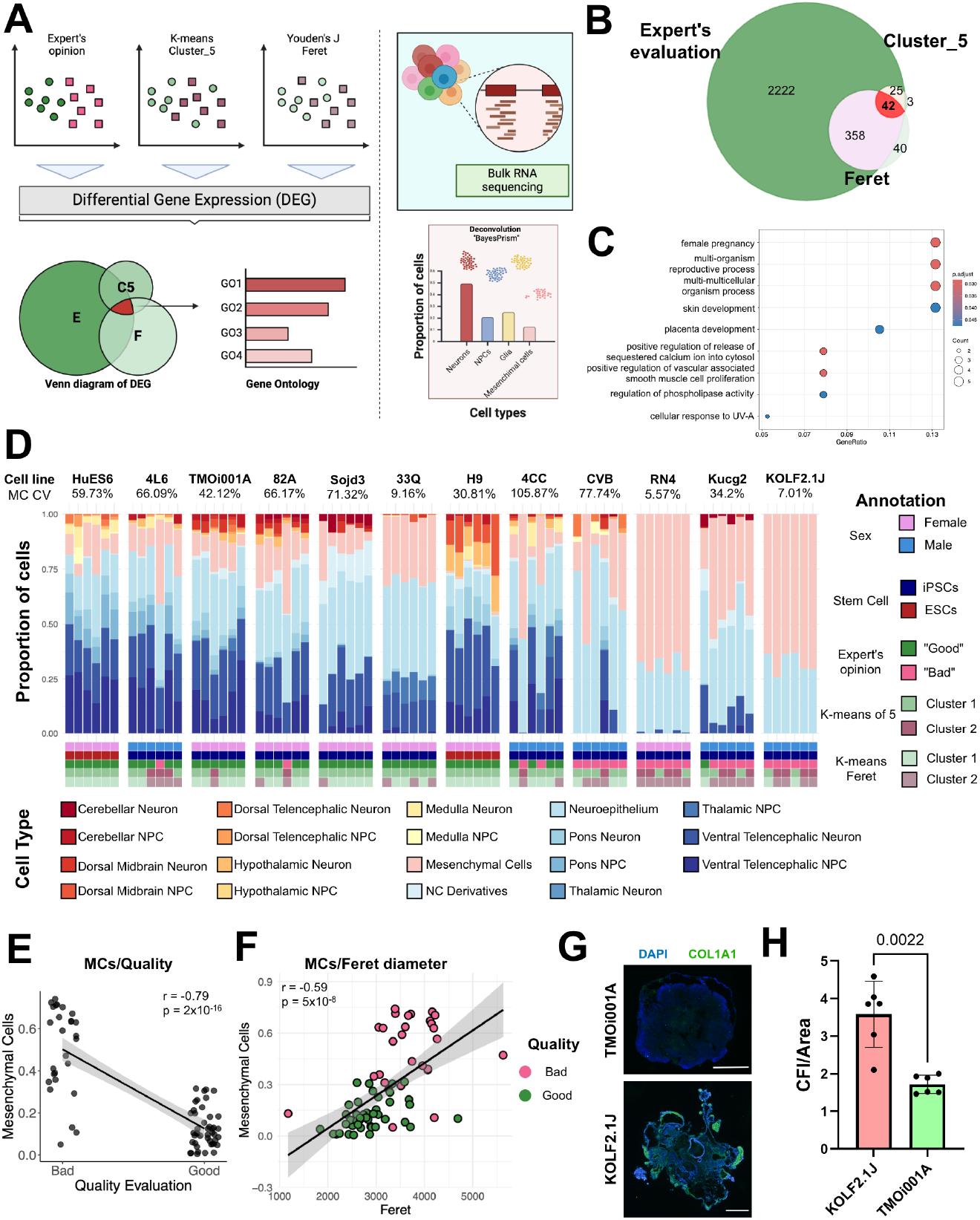
Mesenchymal cells affect brain organoids morphology. **A**. Illustration of the strategy for gene expression analysis. **B**. Venn diagram, indicating the number of differentially expressed genes when organoids are compared by expert evaluation (‘Expert’s evaluation), k-means clustering of five top correlating clusters (‘Cluster_5’) or the determined Feret diameter threshold (‘Feret’). The overlap of all three analyses is depicted in red. **C**. Gene Ontology analysis of differentially expressed genes common across three comparisons. The x-axis represents the Gene Ratio, while the y-axis displays the Gene Ontology (GO) terms associated with Biological Processes. The color scale indicates the adjusted p-value, and the size of the circles reflects the number of genes associated with each GO term. **D**. Stacked bar chart illustrating the cellular composition of organoids calculated with deconvolution analysis. Further variables are color encoded below (sex, PSC type, expert evaluation group, group in k-means clustering, group in Feret diameter thresholding). Annotation coding for cell types is shown at the bottom of the graph. Each stack represents an individual organoid. The mesenchymal cell’s coefficient of variation (MC CV) for each is presented as a percentage under the cell line name. **E**. Scatter plot between the estimated proportion of mesenchymal cells and the expert evaluation of organoid quality. r = −0.79, p = 2×10-16. **F**. Scatter plot of Mesenchymal cells (MC) and Feret diameter. r = 0.59, p = 5×10-8. **G**. IHC of TMOi001A and KOLF2.1J brain organoids stained with COL1A1 (green) and DAPI (blue). **H**. Bar plot with corrected fluorescence intensity (CFI) of COL1A1 signal normalized to the area of the organoid. Each dot represents CFI from one section of one organoid of the analyzed line. Statistical significance was tested with the Mann-Whitney-U test (p = 0.0022) (n = 6). Data represented as mean ± SD.

Finally, we aimed to experimentally validate our finding by staining six organoids of two hPSC lines - one line consistently performing as ‘good’ and the other as ‘bad’ in the expert’s evaluation - with the mesenchymal cell-associated protein COL1A1. The COL1A1 extracellular matrix protein is secreted by mesenchymal cells^29^. Organoids from hPSC lines classifying within the ‘bad’ category exhibited higher COL1A1 staining intensity, thereby confirming our finding (Fig 3G).

These data indicate that the quality of brain organoids can be evaluated using distinct morphological parameters. Notably, these parameters correlate with the presence of mesenchymal cells, as higher-quality organoids tend to contain fewer mesenchymal cells.

## Discussion

Here, we describe the heterogeneity of brain organoids of multiple hPSC lines following unguided differentiation. Using a streamlined data analysis approach, we identify morphological parameters, most prominently the Feret diameter, as a simple-to-estimate surrogate of brain organoid quality. Through bioinformatic deconvolution analysis, we uncover that low-quality brain organoids contain a higher proportion of mesenchymal cells. Hence, our results provide an important cornerstone in the field’s efforts to standardize brain organoid research.

We identified easily measurable morphological features that can be used to objectively assess organoid quality at the early stages of the differentiation process. Among those, the Feret diameter reflected the expert evaluation most precisely. A low Feret diameter indicates a proper brain organoid quality. Area and number of cysts, two other parameters determined in this analysis, have been associated with poor brain organoid quality in earlier studies^19,30^. However, their definite identification may require a certain experience in stem cell research. As brain organoids increase in volume with the number and size of cysts, the Feret diameter reflects this process. Implementing this measurement as a parameter for organoid selection is appealing, as it is a straightforward measurement that can be performed using standard cell culture microscopes. It can be applied by personnel or trainees with minimal experience or even in an automated fashion. Hence, this finding could aid the growing scientific community using brain organoids for specific questions but missing extensive experimental experience in the field.

We aimed to uncover the cellular basis of poor organoid quality by applying state-of-the-art deconvolution analysis of RNA-seq of individual organoids. We found that mesenchymal cells are the main culprit restricting proper brain organoid development. The presence of mesenchymal cells in brain organoids derived from hPSCs has been reported before^5,27,31^, but not further characterized. Mesenchymal stem cells are frequently discussed to have neuroprotective properties in neurological disorders via cell-cell contact or secreted factors and exosomes^32–34^. This cell type was also under clinical investigation in neurodegenerative disorders, e.g., Alzheimer’s disease (Clinical Trial NCT02833792) and amyotrophic lateral sclerosis (Clinical Trial NCT03280056). However, no convincing proof of clinical efficacy leading to FDA or EU approval has been shown so far. In the context of CNS malignancies, the abundance of mesenchymal cells in tumor tissue inversely correlates with patient survival, and mesenchymal cells are thought to promote the proliferation of glioma cancer cells^35^. Hence, the presence of this cell type should caution researchers in disease modeling and preclinical development, as it may significantly confound results. Therefore, pre-screening brain organoids using a defined morphological parameter to obtain organoids with fewer cells of confounding potential can be an asset in therapeutic development.

The reported variability between organoids of the same PSC line and the variability between different PSC lines from healthy individuals underscores the need for careful considerations regarding organoid selection when comparing phenotypes between healthy and diseased conditions. The common practice in the field to choose individual organoids by an expert evaluation will require, at a minimum, a more detailed methodological presentation of this arbitrarily appearing selection process concerning brain organoid quality. Ideally, they should also reflect the scientific community’s consensus on standardized parameters for quality aspects of organoid morphology and cellular composition. When generating human (neural) organoid cell atlases integrating multiple single-cell transcriptomics datasets from various protocols and sources comprising millions of cells into a single comprehensive database, special attention should be given to the number and origin of input hPSC lines and the experimenter’s procedure of brain organoid selection prior to omics analyses.

In line with international guidelines and the stem cell community’s efforts, our findings provide new insights into the cellular basis of brain organoid heterogeneity and an easy-to-use framework for researchers to lower experimental variability.

## Materials and Methods

### Resource availability

#### Lead contact

Information and requests for resources and reagents could be directed to and will be fulfilled by Dr. Florian Krach.

#### Materials availability

Please note that there are restrictions to the availability of UKER iPSC lines due to the vote of the ethics committee.

#### Data and code availability

Data has been deposited at the European Genome-phenome Archive (EGA), which is hosted by the EBI and the CRG, under accession number EGAS50000000659. Further information about EGA can be found at https://ega-archive.org and “The European Genome-phenome Archive of human data consented for biomedical research.

### Ethics Statement

The human cells used in this study were handled in accordance with the principles outlined in the Declaration of Helsinki. Concerning all UKER lines, informed written consent was obtained from the participating individuals to use donor tissue for research purposes. The generation and use of local human iPSC lines were approved by the Institutional Review Board of the University Hospital of Erlangen (Nr. 4120 and 259_17B: Generation of human neuronal models for neurodegenerative diseases). Concerning iPSC lines Kucg2 and Sojd3 (HipSci feeder-free panel (ECACC 77659901)), the MTA was obtained from the Wellcome Trust Sanger Institute for research purposes. The remaining iPSC lines (TMOi001A, KOLF2.1, CVB) are commercially available. The use of human embryonic stem cells for this project was approved by the Central Ethics Committee for Stem Cell Research (121. Approval according to the German Stem Cell Act to Beate Winner).

### Human pluripotent stem cell line maintenance and processing

We used a total of 10 previously published iPSC lines derived from healthy human tissues - UKERi4L6, UKERi4CC, UKERiRN4, UKERi33Q, UKERi82A, TMOi001A, KOLF2.1, Kucg2, Sojd3, CVB, and two human ESC lines – H9 and HuES6 (Table S1).

Cell lines UKERi4L6, UKERi4CC, UKERiRN4, UKERi33Q, and UKERi82A^36,37^ were generated at the University Hospital Erlangen (Erlangen, Germany). TMOi001-A line was bought from Thermo Fisher (Waltham, MA, USA). Kucg2 and Sojd3 lines were purchased from Wellcome Sanger Institute (WTSI) via the European Collection of Authenticated Cell Cultures (ECACC) (Porton Down, UK). The KOLF2.1 cell line^38^ was obtained from the Jackson Laboratory (Bar Harbor, ME, USA). The CVB cell line was bought from the Coriell Institute for Medical Research (Camden, NJ, USA). HuES6 cell line was received from Harvard Stem Cell Institute, and H9 was obtained from WiCell Research Institute (Madison, WI, USA). IPSC lines UKERi4L6, UKERi4CC, UKERiRN4, UKERi33Q, and UKERi82A were generated at the Department of Stem Cell Biology. For iPSC generation, skin biopsies of study participants were obtained. iPSCs were generated from fibroblasts using the CytoTune iPS 2.0 Sendai Reprogramming Kit (Thermo Fisher Scientific, USA) according to the manufacturer’s instructions. All cell lines have been operated in sterile conditions and tested negative for mycoplasma before being included in the experimental procedures. Cells were cultured in human stem cell media mTESR Plus (StemCell Technologies) with mTESR Plus supplements (StemCell Technologies). The medium was changed every other day. Cells were propagated with ReLeSR™ Passaging Reagent (StemCell Technologies, Canada) on 4 mg/ml Geltrex (Gibco, USA) coated polystyrene cell culture plates for growth. Low-pass whole-genome sequencing at a depth of 1.7x was performed for all cell lines using a NovaSeq 6000 (Illumina, USA). Reads were aligned to the human genome reference (hg19), and copy number variation (CNV) analysis was conducted using the DRAGEN CNV pipeline (ver. 3.10).

### Human brain organoid differentiation

To generate brain organoids, iPSCs and ESCs were differentiated using a protocol from Lancaster et al.^19^ with slight adaptations. Briefly, prior to brain organoid differentiation, cells were treated with Accutase (Gibco, USA) for 5 min at 37 °C to dissociate cells and have a single-cell suspension. The cells were resuspended in Organoid Formation Media (OFM) (DMEM/F12 with GlutaMAX, 20% Knockout Serum Replacement, 3% FCS, 1% MEM-NEAA, 50 µM β-Mercaptoethanol (all Gibco, USA) with 50 µM Rock inhibitor and 4 ng/ml FGF2 (PeproTech, USA). Embryoid bodies (EBs) were generated by adding 10,000 cells to each well of a 96 U-bottomed plate, which was then centrifuged at 600 g for 5 minutes and incubated in OFM supplemented with ROCK inhibitor (Enzo Life Sciences, Germany) and FGF2 (PeproTech, USA). On day 3, two-thirds of the media were replaced with fresh OFM without ROCK inhibitor and FGF2 supplements. On day 5, the media was completely changed from OFM to NIM (Neuronal Induction Media) consisting of DMEM/F12 with GlutaMAX, 1% N2, 1% MEM-NEAA, and 1 µg/mL Heparin. The media was changed every other day thereafter. On day 11, each organoid was embedded into a Matrigel (Corning, USA) droplet and placed into a 6-well plate with Cerebral Differentiation Media (CDM) without Vitamin A (50% DMEM/F12 (Gibco, USA), 50% Neurobasal (Gibco, USA), 0.5x N2 (17502048, Gibco), 1x B27 (12587-010, Invitrogen), 1x Pen/Strep (15140-122, Gibco), 0.5x MEM-NEAA (11140-35, Gibco), 0.025% Insulin (I9278, Sigma), 50 µM β-Mercaptoethanol (Carl Roth, Germany)). The organoids were incubated without agitation until day 15 and fed every second day with CDM without Vitamin A. On day 15, the organoid media was changed to CDM with Vitamin A. The 6-well plate was then transferred to an orbital shaker (Laborschüttler Rocker 3D basic, IKA, Germany), rotating at 33 rpm. The media was changed every other day until day 30. On day 30 of differentiation, brain organoids with distinct morphology were chosen for further gene expression analysis using RNA sequencing.

### Bright-field imaging and morphometry analysis

To monitor the growth dynamics of brain organoids, random organoids from each cell line were selected and imaged every five days from day 11 to day 30 using a Zeiss Axio Vert.A1 microscope. For larger organoids, multiple overlapping images were captured and stitched together to create a single composite image for comprehensive visualization. On day 30, a subset of organoids (six per cell line) was selected for RNA sequencing, and these organoids were imaged using a Zeiss Stemi 2000-CS microscope.

For morphological analysis, images of the organoids were processed using ImageJ software (version 2.14.0). Prior to analysis, all images were converted to 8-bit grayscale format to standardize the processing and ensure compatibility with thresholding functions. Automatic thresholding was applied to distinguish organoid structures from the background, followed by manual adjustments to optimize segmentation and accurately define the organoid boundaries. This ensured that the entire structure of each organoid was captured for analysis.

Subsequent morphological measurements were conducted. Area, Min and Max gray values, Shape descriptors, Mean gray value, Feret diameter, Median, and Kurtosis were initially chosen for morphological characteristics of brain organoid images using the Set Measurements function in ImageJ. Later, the following parameters were selected for a better description of the shape of the organoids: Area, Perimeter, Aspect Ratio, Feret Diameter, Roundness, Circularity, and Solidity. These metrics provided quantitative data on organoid size, shape, and structural integrity. Area and perimeter measurements were used to assess the organoid size, while shape descriptors such as aspect ratio, circularity, and roundness offered insights into the structural form of the organoids. Solidity was particularly useful in evaluating the compactness and overall structural integrity of each organoid.

On day 30, cystic structures within the organoids were also evaluated. Cysts were identified based on their morphology and quantified by counting the number of cysts per organoid. Additionally, the total cyst area was measured and expressed as a proportion of the entire organoid area to assess the prevalence of cyst formation across different cell lines. These data, along with other morphological measurements, are presented in the supplemental source data table.

### Flow cytometry (FC)

The pluripotency of hPSCs was evaluated by flow cytometry using the TRA-1-60 marker (Biotec). To assess cellular composition, we quantified PAX6 and MAP2 cells in brain organoids. Brain organoids were dissociated by recovering them from Matrigel using a Cell Recovery Solution (Corning) and then using a papain-based Neural Tissue Dissociation Kit (Miltenyi Biotec) according to the manufacturer’s instructions. Live-dead staining was performed with the LIVE/DEAD™ Fixable Dead Cell Stain Kit (Thermo Fisher Scientific) as per protocol. For intracellular staining, 5×10^5^ cells were fixed with 4% paraformaldehyde for 15 minutes at room temperature. Cells were permeabilized and blocked with 2% FCS and Fc receptor block (BioLegend). They were then incubated with anti-PAX6-APC (1:50, Miltenyi Biotec, 130-123-267) and anti-MAP2-PE (1:100, Merck, FCMAB318PE) antibodies for 15 minutes at 4°C in the dark. Flow cytometry was conducted using a CytoFLEX flow cytometer and analyzed with CytExpert software version 2.4.0.28 (both from Beckman Coulter). Fluorescence compensation was performed using UltraComp eBeads™ Compensation Beads (Invitrogen). For flow cytometry analysis, cells from the samples underwent gating for single cells, followed by gating for viable cells. They were then analyzed for MAP2 and PAX6 positivity. Unstained and fluorescence-minus-one samples were used as controls to establish gating parameters. Three brain organoids per cell line were analyzed for cellular composition via flow cytometry.

### Immunohistochemistry (IHC)

On day 30, organoids were washed twice with PBS and fixed in 4% paraformaldehyde (PFA, Carl Roth) for 1 hour. Post-fixation, they were washed thrice with PBS (10 minutes each) and immersed in 30% sucrose for cryoprotection. Organoids were then embedded in Neg-50™ Frozen Section Medium (Thermo Fisher) on dry ice and stored at −20°C. Organoids were cryosectioned into 30 μm slices using a CryoStar NX70 cryostat (Thermo Fisher) and placed on SuperFrost Plus™ slides (Thermo Fisher). Prior to antigen retrieval, organoid sections were fixed again in 4% PFA for 12 minutes, followed by two washes in PBS. For antigen retrieval, sections were incubated at 70°C for 20 minutes in HistoVT One buffer (1:10 dilution, Thermo Fisher). Sections were blocked for 15 minutes with PBS containing 4% normal donkey serum (NDS, Sigma-Aldrich) and 0.25% Triton X-100 (Sigma-Aldrich), followed by a 2-hour incubation at room temperature with the same blocking buffer. Primary antibodies diluted in antibody solution (PBS, 4% NDS, 0.1% Triton X-100) were applied overnight at 4°C (SOX2: 1:300 (3579S, Cell Signaling Technology); MAP2: 1:500 (M9942, Sigma-Aldrich); COL1A1 (3G3): 1:100 (sc-293182, Santa Cruz)). The next day, sections were washed twice with PBS (5 minutes each) and once with PBS containing 0.5% Triton X-100 (8 minutes). Secondary antibodies and DAPI (1:1000, Sigma-Aldrich) were applied for 2 hours at room temperature. Sections were washed thrice with PBS (5 minutes each) and mounted using Aqua Polymount (Polysciences). Immunofluorescence images were captured using an EVOS M7000 Imaging System (Thermo Fisher) for SOX2 and MAP2 staining and a Zeiss Observer Z1 fluorescence microscope (Zeiss) for COL1A1 staining. To evaluate the signal intensity from the COL1A1 antibodies, we measured the Corrected Fluorescent Intensity *CFI* = *Integrated Density* − (*Area × Background Mean Intensity*)), using the ImageJ software version 2.14.0 and normalized it to the size of the organoid.

### Total RNA isolation, quality control, library preparation, and sequencing

RNA was extracted using the RNeasy kit (Qiagen, Germany) according to the manufacturer’s instructions. RNA concentrations were measured using a NanoDrop NP80 (Implen, Germany). A total of 1000 ng per sample was sent for RNA sequencing to Azenta Life Sciences (Genewiz Leipzig, Germany) for sequencing library preparation and 150 bp paired-end sequencing with Poly-A selection. 72 samples were sent and sequenced at a depth of >20 million reads in each sample. RNA samples were quantified using a Qubit 4.0 Fluorometer (Life Technologies, Carlsbad, USA), and RNA integrity was checked with an RNA Kit on an Agilent 5300 Fragment Analyzer (Agilent Technologies, Palo Alto, USA). The ERCC RNA Spike-In Mix kit (Thermo Fisher Scientific, USA) was added to normalized total RNA prior to library preparation following the manufacturer’s protocol. RNA sequencing libraries were prepared using the NEBNext Ultra II RNA Library Prep Kit for Illumina following the manufacturer’s instructions (New England Biolabs, USA). mRNAs were first enriched with Oligo(dT) beads. The samples were sequenced using a 2×150 Pair-End configuration (ver. 1.5) on an Illumina NovaSeq 6000. Image analysis and base calling were conducted by the NovaSeq Control Software (ver. 1.7). Raw sequence data was converted into .fastq files using the bcl2fastq program version 2.20.

### RNA seq analysis

To assess which gene expression affects organoid morphology and quality, we performed RNA sequencing of 72 organoids derived from 12 different stem cell lines. FastQC was used for the quality control of raw sequences (Andrews, S. (2010). FastQC: A Quality Control Tool for High Throughput Sequence Data. Available online at: http://www.bioinformatics.babraham.ac.uk/projects/fastqc/). The raw sequence reads were aligned to the human reference genome GRCh38 using the STAR aligner (ver. 2.7.11)^39^. Gene expression levels were quantified using the featureCounts tool as part of the Rsubread package (ver. 2.16.1)^40^. FeatureCounts assigns reads to genomic features, such as exons and genes, to produce count data representing the abundance of each gene in the samples. Three distinct DE analyses were performed for comparisons across the different conditions using the DESeq2 R package (ver. 1.42.1)^41^. These three conditions compared “Good” and “Bad” quality organoid samples according to the Expert’s evaluation (Expert), k-means clustering based on five morphological parameters (Cluster_5), and Feret diameter (Feret) threshold 3050 mcm. To adjust the differences between different cell lines, the design formula included Cell lines as a covariate (design: ∼ Condition + Cell_lines). Lowly expressed genes were excluded using the quartile method by retaining only those with a mean expression above the first quartile (25th percentile) across all samples prior to differential gene expression analysis. For each condition (Expert, Cluster_5, Feret), differential expression analysis was performed using the DESeq2 pipeline. Only genes with an adjusted p-value (padj) less than 0.05 and a |log2FC| > 1 are considered significantly differentially expressed. PCA was performed on the VST-transformed data for each condition to visualize the variance in gene expression between the different experimental groups. PCA plots were generated using the ggplot2 R package (ver. 3.5.1) for aesthetic adjustments, including color-coding of conditions (‘Bad’, ‘Good’) using a custom color palette. ENSEMBL gene identifiers were mapped to gene symbols using the org.Hs.eg.db annotation package (ver. 3.18.0). Gene Ontology (GO) enrichment analysis was conducted using the clusterprofiler (ver. 4.10.1)^42^ and enrichplot (ver. 1.22.0) (https://github.com/YuLab-SMU/enrichplot) packages. Gene Ontology enrichment analysis was performed on the significantly differentially expressed genes (padj < 0.05; |log2FC| > 1) for each condition using the clusterProfiler package (ver. 4.10.1). Venn diagram was made with eulerr (ver. 7.0.2) package. As background genes for GO enrichment analysis, we used genes included in the differential gene expression analysis. Differential gene expression and GO analysis and visualization have been performed in RStudio software (ver. 2023.09.1+494).

### BayesPrism deconvolution assay

To estimate approximate cell type proportions in the organoids, we performed a deconvolution analysis using BayesPrism (ver. 2.0)^26^. As a reference, we used samples from organoids produced using the Lancaster protocol between day 20 and day 50 from the Human Neural Organoid Cell Atlas^27^. We used the level 3 annotation, removing labels with fewer than 100 cells in the selected samples, leaving 39898 cells with 19 labels. Protein-coding genes excluding mitochondrial, ribosomal, sex chromosome and MALAT1 genes were used to perform the deconvolution.

### Correlation analysis

We assessed the correlation between the proportion of mesenchymal cells (MC) and the Feret diameter of organoids using Pearson’s correlation coefficient. We calculated the coefficient (r) and its associated p-value to evaluate the strength and significance of this correlation. For analysis and visualization, we used R (version 4.0.2) (R Core Team (2023)) along with the tidyverse (version 2.0.0) package^43^.

### K-means clustering analysis

K-means clustering was utilized to objectively categorize organoid groups based on their morphological data using R version 4.0.2 (R Core Team (2023)), along with the stats, tidyverse (version 2.0.0)^43^, and factoextra (version 1.0.7) packages. Clustering was performed using five morphological parameters (Feret diameter, Area, Cysts Amount, Cysts Area, and Perimeter), chosen after quality correlation analysis. The optimal number of clusters (k) was determined using the elbow method (Supplementary Figure S1). The total within-cluster sum of squares (WSS) was calculated for k values ranging from 1 to 15. The ‘elbow’ point in the plot of WSS against k indicated the optimal number of clusters (limiting k to >1), resulting in k = 2 for both analyses. Data was normalized using Z-score normalization. K-means clustering was performed with 20 random starts to increase the likelihood of finding the global optimum. The random seed was set to 111 for reproducibility in both clustering analyses.

### Youden’s J Statistic

We used Youden’s J statistic to evaluate nine morphological parameters (Feret diameter, Area, Cyst Amount, Cyst Area, Perimeter, Circularity, Aspect Ratio, Roundness, and Solidity) for their ability to distinguish between ‘good’ and ‘bad’ organoids. We used R version 4.0.2 along with the stats, cutpointr (ver. 1.1.2)^44^, tidyverse (ver. 2.0.0)^43^, and pROC (ver. 1.18.5)^45^ packages. For each parameter, the optimal cutoff point was determined using the cutpointr function. Youden’s index (J) was calculated using the formula: *J* = *Sensitivity* + *Specificity* − 1. Sensitivity and specificity were computed across all cutoff values for each parameter, with the optimal cutoff identified where Youden’s Index was maximized, indicating the best balance.

### Statistical analysis

Sample variance was quantified according to the coefficient of variation: 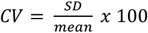. Statistical analyses were performed using RStudio (ver. 2023.09.1+494) and Prism (ver. 10.2.0). The figures’ legends include details for statistical analyses, including replicate numbers. For pair-wise comparisons, a significance was assumed when the p-value was below 0.05.

## Acknowledgments

This study was supported by the Bavarian Ministry of Science and the Arts in the framework of the Bavarian Research Consortium ‘Interaction of Human Brain Cells’ (ForInter) network. Additional support came from the Else Kröner-Fresenius-Stiftung (2024_EKEA.50 (F.K.)), German Research Foundation, DFG (WI 3567/2-1 (BW); GRK2162/ 270949263 (B.W. and J.W.), CRU5024/ 505539112 WI 3567/4-1 (B.W.) and WI 1620/4-1 (JW)), the TreatHSP consortium (BMBF 01GM1905B, 01GM2209B to BW, MR and JW) and the Interdisziplinäres Zentrum für Klinische Forschung (IZKF) (Erstantragsteller project J-112 (D.K.)); ELAN-Fond P153 (F.K.). The authors thank Holger Wend for excellent technical support.

Graphical illustrations were made with BioRender.com.

## Author Contributions

Conceptualization - D.K., T.B., B.W., F.K., M.K., S.F.; Methodology - D.K., M.F., S.P., L.Z., S.F., S.P., M.F., P.G.; Investigation - D.K., M.F. N.Z., F.F., N.N., M.B.; Formal analysis - D.K., N.Z., F.K.; Visualization - D.K., N.Z., L.Z.; Data curation - D.K., N.Z.; Original draft preparation - D.K., F.K., B.W.; Writing - Review and Editing - D.K., F.K., N.Z., B.W., F.T.; Project administration - D.K., F.K., B.W.; Funding acquisition - B.W., M.R., J.W., C.G., D.K., F.K. All authors critically revised the manuscript and approved the submitted version.

