## Supplementary Tables and Figures for "Deciphering Brain Organoids Heterogeneity by Identifying Key Quality Determinants"

### Morphology and Mesenchymal Cell Proportion Define the Quality of Brain Organoids

### Table S1. Cell lines used in the study

| Cell line | Short naming | Cell type | Gender | Passage | |
|---|---|---|---|---|---|
| WA09, WAe009-A | H9 | hES | Female | p41 | |
| HVRDe006-A | HuES6 | hES | Female | p35 | |
| KOLF2.1J | KOLF | iPSC | Male | p+4* | * - passage from the moment the line was obtained |
| WTSIi013-A | Kucg2 | iPSC | Male | p38 | |
| HPSI0314i-sojd_3 | Sojd3 | iPSC | Female | p33 | |
| CV-hIPS-B | CVB | iPSC | Male | p38 | |
| TMOi001A | Thermo | iPSC | Female | p26 | |
| UKERi4CC-S1-015 | 4CC | iPSC | Male | p16 | |
| UKERiRN4-S1-009 | RN4 | iPSC | Female | p12 | |
| UKERi4L6-S1-027 | 4L6 | iPSC | Male | p20 | |
| UKERi33Q-S1-101 | 33Q | iPSC | Female | p12 | |
| UKERi82A-S1-002 | 82A | iPSC | Female | p15 | |

### Table S2. Youden's J Statistics for brain organoid morphological parameters

| Parameter | Optimal cutoff | Youden's Index | Sensitivity (%) | Specificity (%) |
|---|---|---|---|---|
| **Area** | 5695105.348 | 0.67 | 77% | 88.9% |
| **Perimeter** | 13400.056 | 0.65 | 96.3% | 68.9% |
| **Circularity** | 0.298 | 0.52 | 77.8% | 74.1% |
| **Feret** | 3050.356 | 0.68 | 92.6% | 75.6% |
| **Aspect Ratio** | 1.217 | 0.33 | 77.8% | 55.6% |
| **Roundness** | 0.824 | 0.33 | 55.6% | 77.8% |
| **Solidity** | 0.909 | 0.53 | 82.2% | 70.4% |
| **Cysts.Area** | 0.0895 | 0.67 | 96.3% | 71.1% |
| **Cysts.Amount** | 2 | 0.53 | 81.5% | 71.1% |

### Figure S1. K-means clustering. Scree plot

**A**

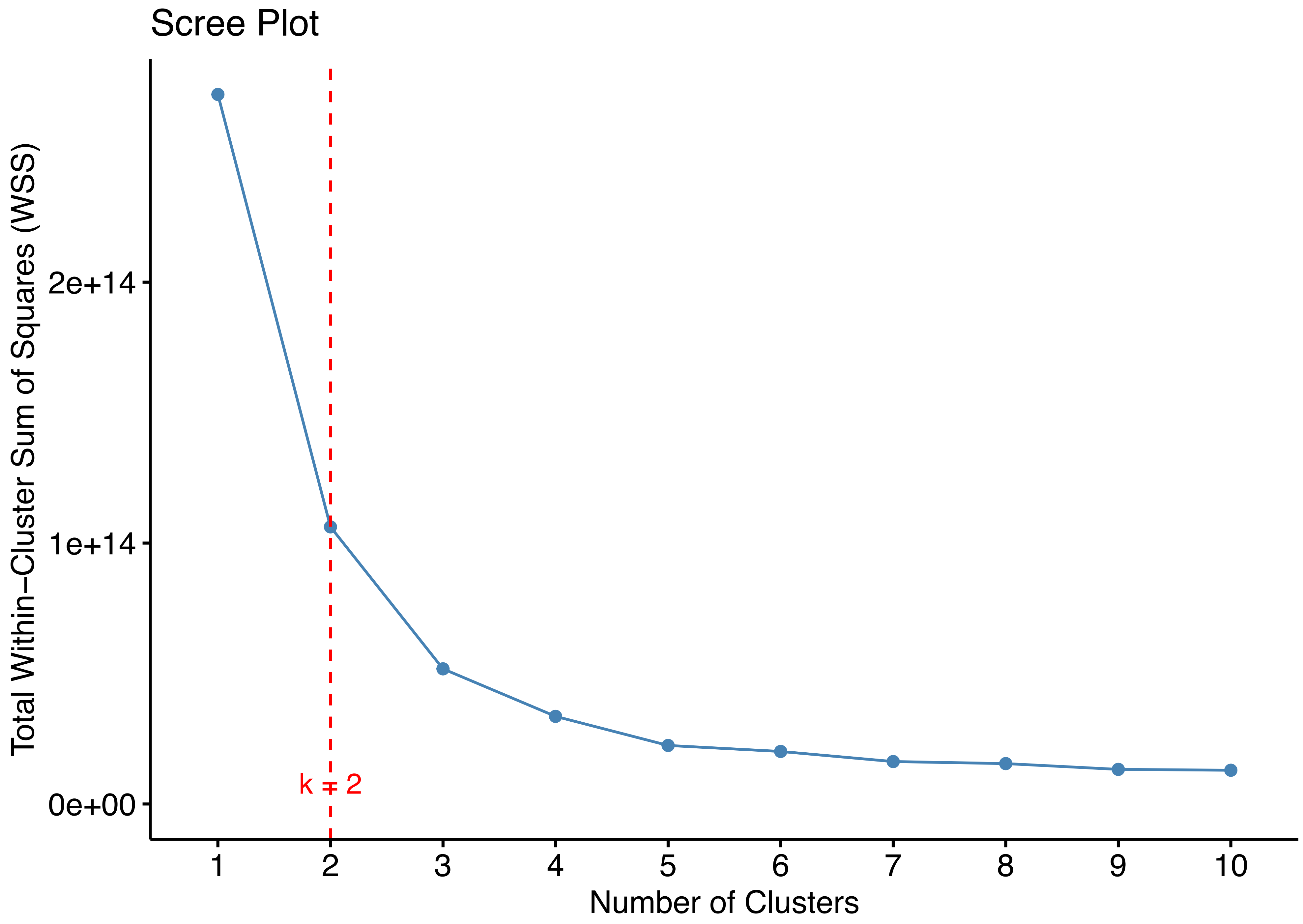


**Figure S2. The k-means elbow method is used to determine the optimal number of clusters.** The x-axis represents the number of clusters, ranging from 1 to 10. The y-axis displays the Total Within-Cluster Sum of Squares (WSS). A red dashed vertical line marks the position where k equals 2 - the suggested optimal number of clusters based on the elbow method.

### Figure S2. Morphology-based PCA plot of the d30 brain organoids

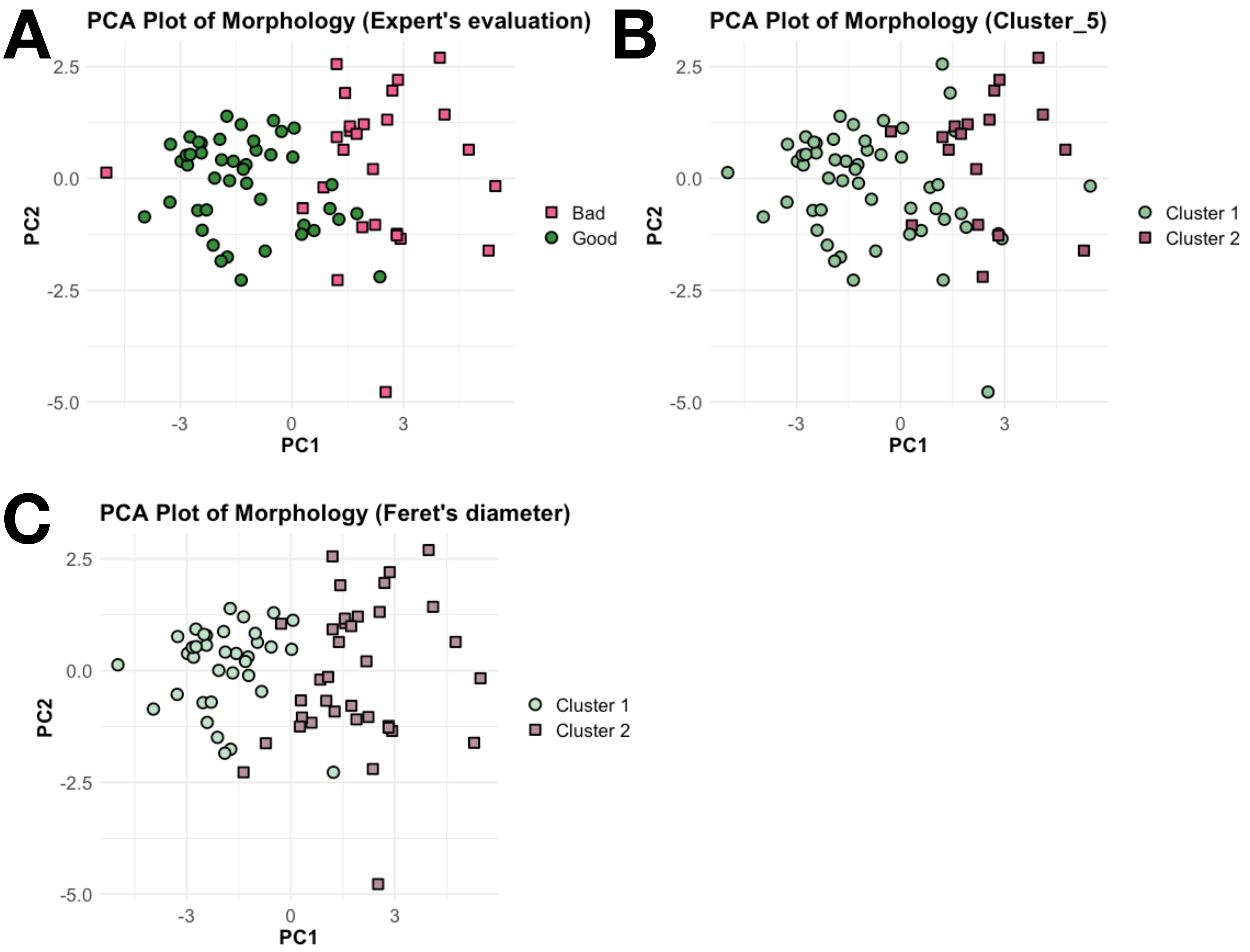


**Figure S3. Morphology-based PCA plot of day 30 (d30) brain organoids.**

Principal Component Analysis (PCA) plot illustrating the morphology of d30 brain organoids, with samples color-coded according to various quality assessment methods. **A.** Samples are color-coded based on Expert evaluation of organoid quality: green represents 'Good' organoids, and pink represents 'Bad' organoids. **B.** Samples are color-coded according to k-means clustering results, using five morphological parameters (Feret, Area, Cyst Amount, Cyst Area, and Perimeter). Cluster 1 corresponds to the 'Good' organoid group, while Cluster 2 aligns with the 'Bad' organoid group, as per the expert evaluation. **C.** Samples are color-coded based on Youden's J statistic for the optimal cutoff of Feret diameter. Cluster 1 represents organoids classified as 'Good' (less than 3050 µm in diameter), and Cluster 2 represents organoids classified as 'Bad' (more than 3050 µm in diameter), consistent with expert evaluation.

The variance explained by Principal Component 1 (PC1) is 55.58%, and by Principal Component 2 (PC2) is 17.88%.

### Figure S3. WebCSEA - Web-based Cell-type Specific Enrichment Analysis of Genes for Cell type determination

A

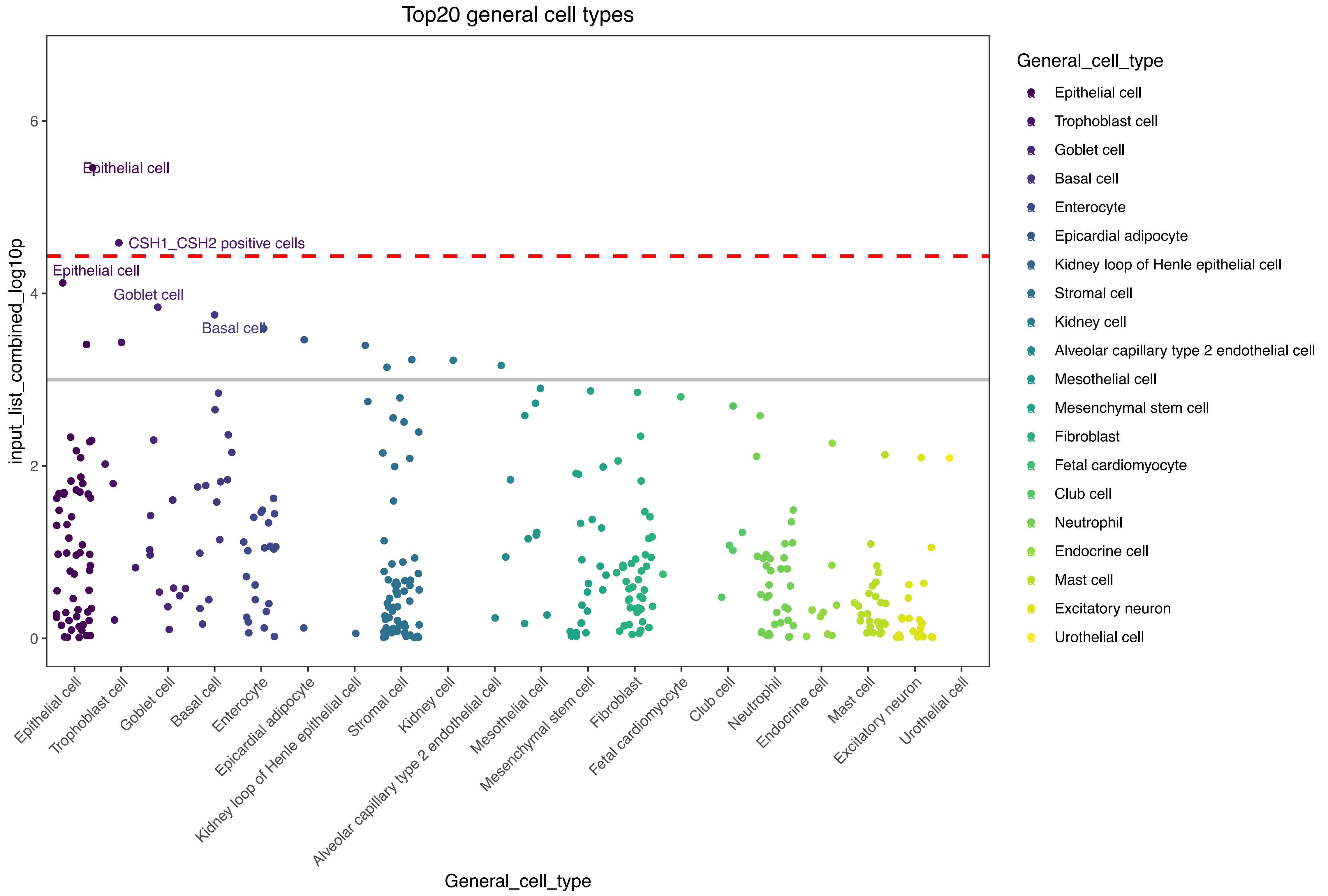


**Figure S4. WebCSEA - Web-based Cell-type Specific Enrichment Analysis of Genes for Cell Type Determination. A.** Jitter plots of –log10 raw (left) P-values among 1355 tissue-cell types by top 20 enriched general cell types, defined by the WebCSEA, based on the list of common differentially expressed genes from the three brain organoid evaluation approaches. The red dashed line indicates the Bonferroni-corrected significance ($P = 3.69 \times 10^{-5}$) by 1355 tissue-cell types. The grey solid line indicates the nominal significance ($P = 1 \times 10^{-3}$).
